# Single-Assay Characterization of Ternary Complex Assembly and Activity in Targeted Protein Degradation

**DOI:** 10.1101/2025.08.20.671298

**Authors:** Corey H. Yu, Vi Dougherty, Dongwen Lv, Pirouz Ebadi, Shikha Dubey, Digant Nayak, Anindita Nayak, Daohong Zhou, Shaun K. Olsen, Dmitri N. Ivanov

## Abstract

Targeted protein degradation (TPD) is a rapidly advancing therapeutic strategy that selectively eliminates disease-associated proteins by co-opting the cell’s protein degradation machinery. Covalent modification of proteins with ubiquitin is a critical event in TPD, yet the analytical tools for quantifying the ubiquitination kinetics have been limited. Here, we present a real-time, high-throughput fluorescent assay utilizing purified, FRET-active E2-Ub conjugates to monitor ubiquitin transfer. This assay is highly versatile, requiring no engineering of the target protein or ligase, thereby accelerating assay development and minimizing the risk of artifacts. The single-step, single-turnover nature of the monitored reaction enables rigorous and quantitative analysis of ubiquitination kinetics. We show that this assay can be used to measure key degrader characteristics such as degrader affinity for the target protein, degrader affinity for the ligase, affinity of ternary complex assembly, and catalytic efficiency of the ternary complex. The high sensitivity and accuracy of this comprehensive, single-assay approach to ternary complex characterization will empower the discovery and optimization of heterobifunctional degraders and molecular glues.

## INTRODUCTION

Protein ubiquitination is a fundamental regulatory mechanism implicated in virtually every aspect of eukaryotic biology^1^. Its functional versatility arises largely from the intricate regulation and substrate specificity of E3 ubiquitin ligases, a diverse class of enzymes that catalyze ubiquitin transfer from high-energy E2–Ub thioester conjugates onto target proteins^2^. The growing interest in ubiquitin ligases as therapeutic targets has spurred efforts to develop improved tools for monitoring E3-catalyzed ubiquitin transfer^3–5^, yet broadly applicable and robust analytical methods remain limited. This need is particularly pressing in the emerging field of TPD—a promising therapeutic strategy that eliminates pathogenic proteins by using small molecules—heterobifunctional degraders (or proteolysis targeting chimeras: PROTACs) and molecular glues—to recruit these target proteins to cellular ubiquitin ligases, thereby promoting their ubiquitination and subsequent degradation^6^.

The thioester-linked covalent conjugate of ubiquitin with an E2 ubiquitin-conjugating enzyme is a conserved, high-energy intermediate in protein ubiquitination and related post-translational modifications^7–10^. The universality of this intermediate, combined with the stoichiometric consumption of one E2–Ub conjugate molecule per E3-catalyzed turnover, makes quantifying E2–Ub depletion an attractive strategy for monitoring E3 enzyme kinetics. To enable such an analytical assay, we synthesized and purified FRET-active E2– Ub thioester conjugates for a selection of E2 enzymes^11^. The proximity of donor and acceptor fluorescent dyes within the FRET-active conjugate generates a strong FRET signal that gradually decays as the conjugate is consumed during E3-catalyzed ubiquitin transfer^11,12^. Robust assay performance depends on efficient fluorescent labeling of ubiquitin and E2, which we achieved using sortase-catalyzed transpeptidation^13^.

FRET-active conjugates for many different E2 conjugating enzymes have now been synthesized and are commercially available (*Methods*). In this study, we sought to evaluate the utility of these reagents for the discovery and optimization of targeted protein degraders. We demonstrate that the FRET-based assay enables quantitative characterization of E3 ligase kinetics in the presence of molecular glues and heterobifunctional degraders. Our findings establish a versatile, accessible tool for accelerating the development of targeted protein degraders and deepening mechanistic understanding of ubiquitin ligase function.

## RESULTS

### FRET assay of E3 activity

The performance of the FRET assay was first evaluated using a simple two-component ubiquitin transfer reaction catalyzed by an engineered TRIM5α ubiquitin ligase. TRIM5α undergoes initial N-terminal monoubiquitination, followed by the extension of a K63-linked polyubiquitin chain^11,14^. The extension step of this reaction, mediated by the heterodimeric E2, UBE2N/UBE2V2, was monitored by tracking the FRET signal over time (Fig. 1A). Raw fluorescence intensity signals detected at the donor and acceptor emission wavelengths exhibited significant variability due to pipetting errors, air bubbles in microplate wells, and other sample-handling inconsistencies (Fig. 1B). However, this variability was largely eliminated when the normalized FRET signal was calculated as the ratio of acceptor to donor signal intensities. These results demonstrate the robustness of the FRET assay against non-FRET sources of experimental noise.

**Figure 1.**
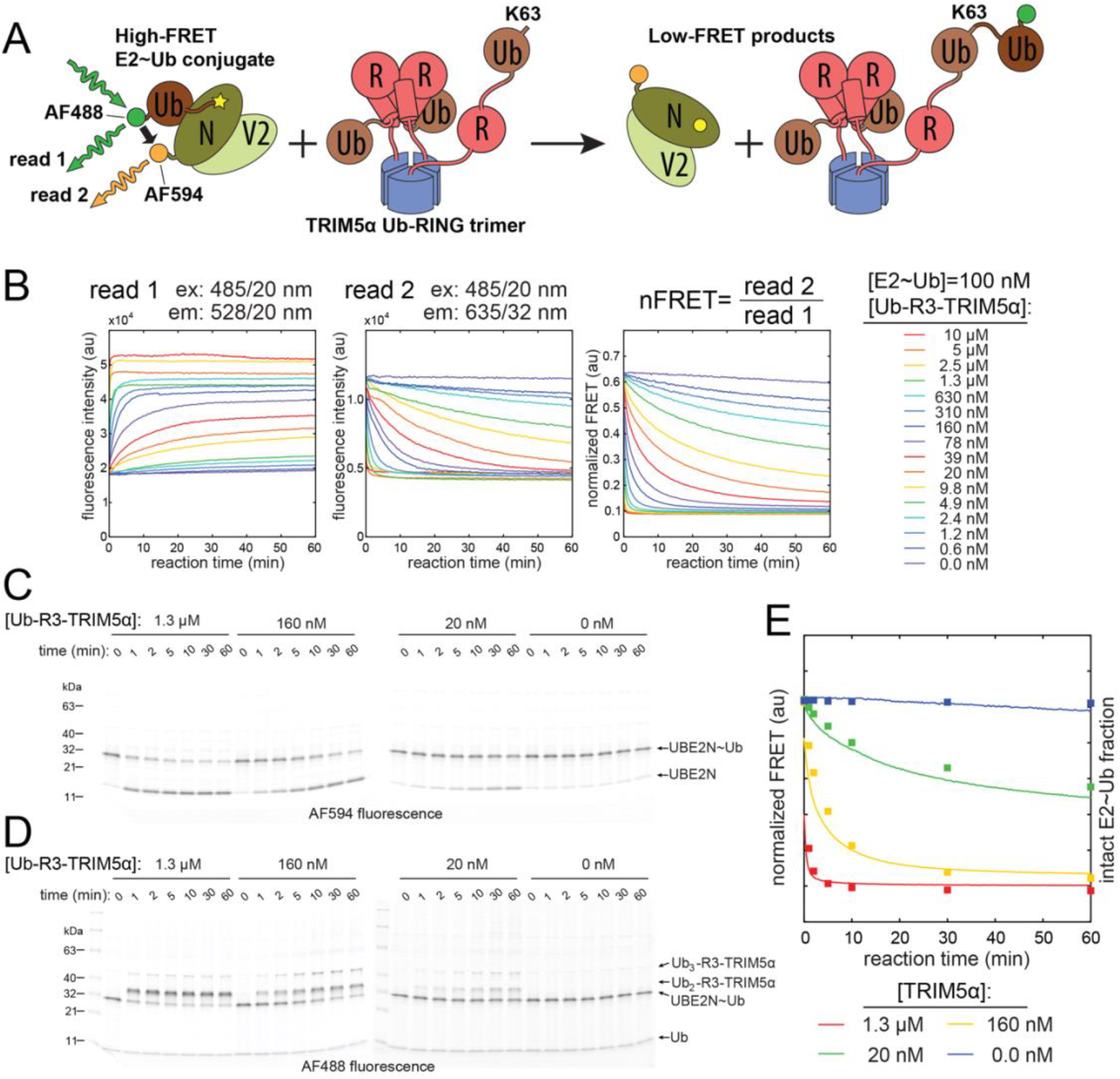
FRET-active E2–Ub conjugates enable real-time monitoring of E3-catalyzed ubiquitin transfer. (A) Schematic of the FRET-based assay using a thioester conjugate of AF488-labeled ubiquitin and AF594-labeled UBE2N. (B) Reaction progress curves recorded by monitoring donor (read 1) and acceptor (read 2) fluorescence emission. Normalized FRET is calculated as the ratio of acceptor to donor fluorescence intensity at each time point. This normalization suppresses non-FRET-related variability due to pipetting or optical artifacts. (C,D) SDS-PAGE analysis of quenched reactions imaged for AF594 (C) and AF488 (D) fluorescence. (E) Overlay of normalized FRET decay curves (lines) and quantification of intact E2–Ub from SDS-PAGE (squares), confirming that FRET signal decay accurately reflects E2–Ub consumption.

To confirm that the decay of the FRET signal results from the dissociation of the FRET-active E2-Ub thioester conjugate, the reaction was quenched at different time points, and its products were analyzed by SDS-PAGE (Fig. 1C,D). Quantification of acceptor (AF594) fluorescence associated with E2-Ub and E2 bands on SDS-PAGE showed excellent correlation with FRET data, confirming that the FRET readout accurately reflects the concentration of the intact E2-Ub conjugate remaining in the reaction (Fig. 1C,E).

Imaging of the same SDS-PAGE gel for donor fluorescence reveals the fate of the AF488-labeled ubiquitin transferred from the E2-Ub conjugate (Fig. 1D). This analysis shows that when the concentration of the E2-Ub conjugate exceeds that of the E3 ligase, the reaction proceeds predominantly in a single-turnover regime, with only one fluorescent ubiquitin transferred onto the Ub-R3-T5α construct. The ability to monitor protein ubiquitination kinetics using a single-step, single-turnover assay enables rigorous and quantitative analysis of reaction kinetics, as described in this study.

### Degrader-dependent enhancement of E3-catalyzed ubiquitin transfer

To explore the utility of the FRET assay for studies of TPD, we selected a well-characterized model system of BRD4 degradation by dBET1 and related compounds that recruit BRD4 to the CRL4^CRBN^ E3 ubiquitin ligase complex^15,16^ (Fig. 2A). The DDB1-cereblon (CRBN) adapter module and the neddylated NEDD8-CUL4A-Rbx1 catalytic module of the CRL4^CRBN^ were generated separately as described in *Methods* and mixed to reconstitute the active CRL4^CRBN^ ligase complex. The FRET-active E2-Ub thioester conjugate used in these experiments comprised UBE2D3 labeled with the AF594 acceptor fluorophore and ubiquitin labeled with fluoresceine donor dye. The BD1-BD2 construct (aa 44-460) of human BRD4 was used as ubiquitination substrate in these experiments.

**Figure 2.**
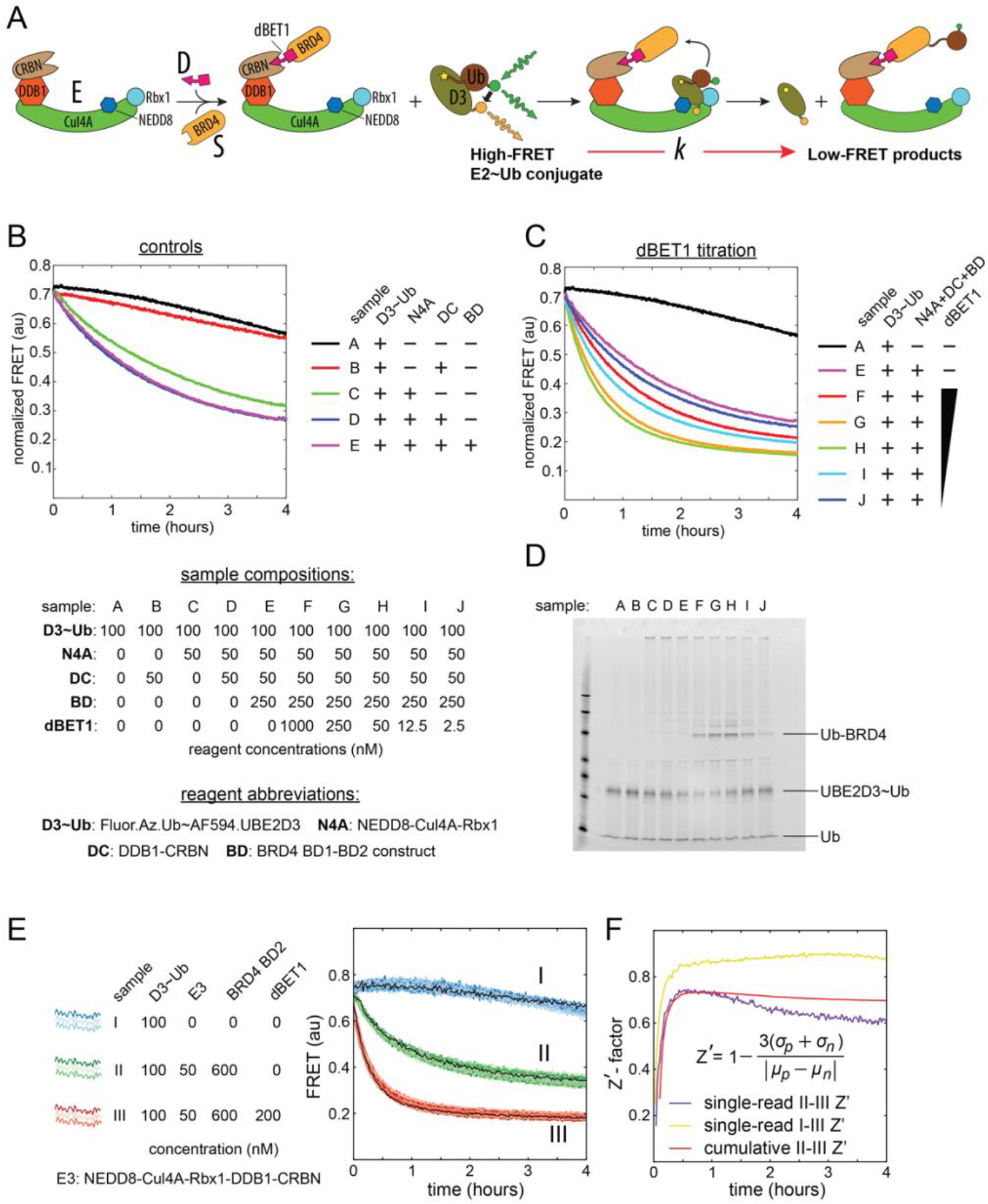
Monitoring degrader-dependent ubiquitin transfer using FRET detection. (A) Schematic of the FRET assay used to monitor dBET1-induced ubiquitination of BRD4 by the CRL4^CRBN^ ligase. (B) Control reactions reveal background activity of neddylated Cul4A–Rbx1 that is independent of both substrate and degrader. (C) Degrader-dependent enhancement of FRET decay shows the characteristic hook effect of heterobifunctional degraders. (D) SDS-PAGE analysis of reaction products at the 1-hour timepoint. Gel imaged for fluorescein fluorescence using a Typhoon scanner. (E) FRET progress curves from 120 replicates for each of three sample compositions (I, II, III), dispensed into a 384-well plate using automated liquid handling. (F) Z′-factors calculated over time for individual reads and cumulative datasets.

Control reactions performed without dBET1 revealed that a significant enhancement of FRET decay occurred in the presence of NEDD8-CUL4A-Rbx1 alone (Fig. 2B). This background activity was only slightly increased by the addition of DDB1-CRBN and was completely independent of the BRD4 substrate. However, when dBET1 was titrated into the reaction mixture containing all components, a clear degrader-dependent acceleration of FRET decay was observed (Fig. 2C). This enhancement increased as dBET1 concentration approached that of the E3 ligase, then declined when dBET1 exceeded BRD4 substrate levels, exhibiting a characteristic “hook effect”^17^. This dependence of FRET decay on degrader concentration suggests that the rate of ubiquitin transfer can serve as a quantitative readout of ternary complex assembly.

To gain further insight, we analyzed the reaction products formed one hour after adding the FRET-active E2-Ub conjugate to the assembled ternary complex using SDS-PAGE (Fig. 2D). Fluorescence imaging of the SDS-PAGE gel revealed that the background activity of NEDD8-CUL4A-Rbx1 alone is highly processive, as indicated by the high molecular weight of the predominant fluorescent band. In contrast, the primary product of the degrader-dependent reaction was monoubiquitylated BRD4 (Fig. 2D).

This result highlights a key distinction between our method, which utilizes purified UBE2D3-Ub conjugates, and traditional in vitro ubiquitination assays, where E1-catalyzed E2-Ub synthesis and E3-catalyzed E2-Ub consumption occur concurrently within the same reaction^18^. In conventional assays with UBE2D3, polyubiquitinated substrate species are typically observed. However, in reactions with preformed UBE2D3-Ub conjugates, the monoubiquitylated product predominates. This observation aligns with previous findings that Cullin-RING ligases (CRLs) utilize UBE2D isoforms for initial monoubiquitination (priming) of the substrate, whereas polyubiquitin chain extension is primarily mediated by UBE2R^19^.

The underlying mechanism of the highly processive background activity observed with NEDD8-CUL4A-Rbx1 and UBE2D3, which contrasts with the degrader-mediated BRD4 monoubiquitination, remains to be elucidated. One possibility is that additional ubiquitination factors or substrates copurify with the CUL4A-Rbx1 complex, which was affinity purified after co-expression of CUL4A and Rbx1 in insect cells (*Methods*).

### Assay Performance in the High-Throughput Screening Format

To evaluate the performance of the FRET assay in a high-throughput format, we measured the Z’-factor, a widely accepted metric for assessing assay robustness and reliability^20^. Z’-factor measurements were conducted in a 384-well plate using automated reagent dispensing, as described in the *Methods* section. The plate contained 120 replicates for each of three distinct sample compositions (I, II, and III), with a final volume of 6 μL per well. Sample III, which contained all components necessary for degrader-dependent substrate ubiquitination, served as the positive control for Z’-factor calculations. Samples I and II were used as negative controls, monitoring E3-independent and degrader-independent consumption of the E2-Ub conjugate, respectively.

The E2-Ub conjugate was the final component dispensed into the plate, initiating the ubiquitination reaction. The reaction was monitored for 4 hours, with FRET measurements taken every 75 seconds across the entire 384-well plate. When the Z’-factor was calculated for each FRET measurement—using Sample II as the negative control and Sample III as the positive control—it peaked at 0.73 approximately 30 minutes after the reaction started. The Z’-factor gradually declined after the initial 30 minutes but remained above 0.6 for the entire 4-hour reaction duration. These results strongly support the assay’s suitability for high-throughput discovery and characterization of targeted protein degraders. Moreover, the assay may be even more effective for HTS identification of E3 inhibitors, as Z’-values calculated between Samples I and III exceeded 0.8.

In addition to single-read Z’-values, cumulative Z’ can be calculated by incorporating data from all reads between t = 0 and a given time point for each sample (*Methods*). This approach greatly reduces the contribution of signal detection noise, while maintaining the contribution of sample preparation variability. The cumulative Z’-factor closely matched individual read Z’-values, suggesting that sample preparation variability (e.g., E3 dispensing errors) is the primary limiting factor of assay performance. This also indicates that reagent concentrations, sample volumes, or data acquisition times could be further reduced without compromising assay performance.

### Time-dependent inactivation model of E3 activity

The accuracy and sensitivity of the FRET assay present an opportunity for rigorous quantitative analysis of E3 ubiquitin ligase kinetics, addressing a long-standing challenge in the field. For example, the strong dependence of FRET decay on degrader concentration suggests that both the concentration and activity of the ternary complex can, in principle, be derived from reaction rate measurements (Fig. 4A).

**Figure 3.**
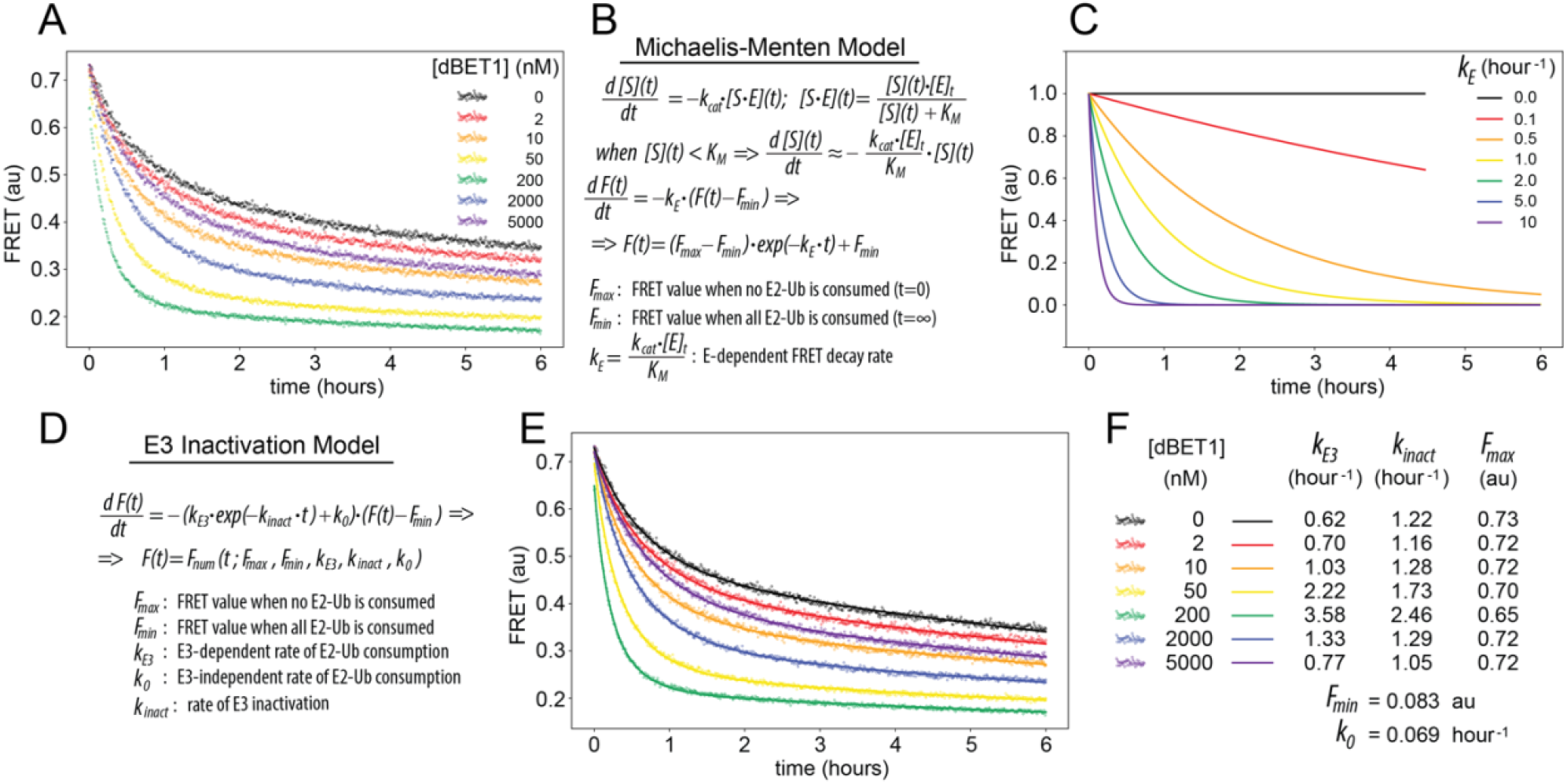
Kinetic modeling of FRET decay rates. (A) FRET decay progress curves from a dBET1 titration series monitoring BRD4 ubiquitination by CRL4^CRBN^. (B) In Michaelis–Menten kinetics, substrate depletion below *K*_*m*_ is well approximated by a simple exponential decay. (C) Enzyme titration series under Michaelis–Menten conditions yield a family of exponential decay curves bounded by the same *F*_*max*_ and *F*_*min*_ values. (D) The differential equation describing time-dependent E3 ligase inactivation can be solved numerically to yield a theoretical substrate depletion curve. (E,F) Global least-squares fitting across the titration series determines best-fit model parameters, yielding excellent agreement between theory and experiment.

**Figure 4.**
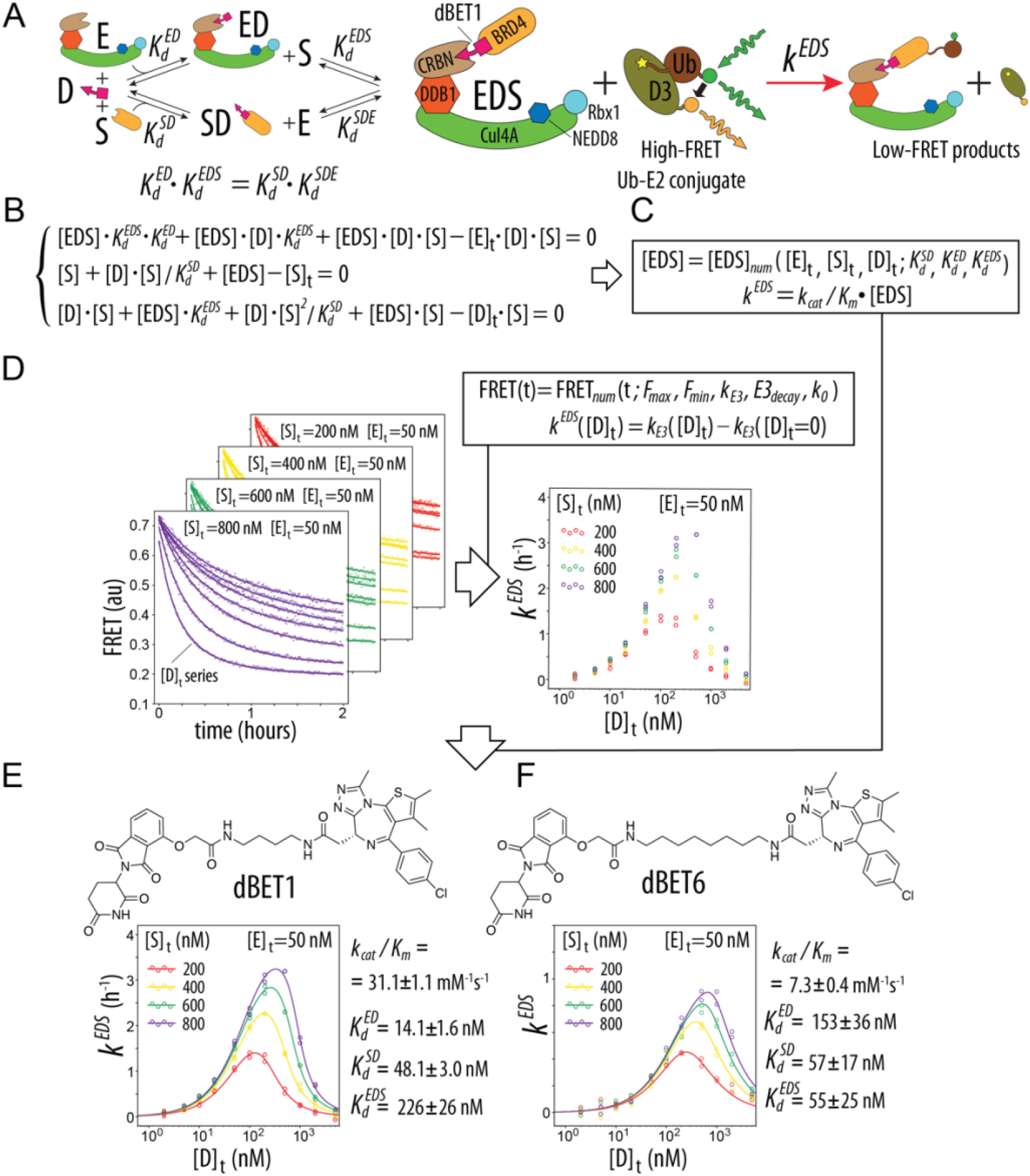
Quantitative analysis of ternary complex assembly using FRET-based ubiquitin transfer assay. (A) Schematic of the reaction and binding equilibria underlying ternary complex formation by heterobifunctional degraders. (B) System of three mass conservation equations describing the equilibria depicted in panel A. (C) Ternary complex concentration [*EDS*] and its associated FRET decay rate *k*^*EDS*^ are calculated by numerically solving the system of equations in panel B. (D) Experimental *k*^*EDS*^ values obtained from degrader titration series at multiple substrate concentrations. The BD2 (aa 333-460) construct of BRD4 was used as ubiquitination substrate. (E,F) Nonlinear least-squares fitting of experimental (D) and modeled (C) *k*^*EDS*^ values yields four parameters describing ternary complex assembly and activity: 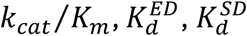 and 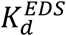.

Under Michaelis-Menten kinetics, the rate of substrate depletion is proportional to the concentration of the Michaelis complex [*S* · *E*]. When the substrate concentration is significantly lower than *K*_*m*_—an assumption that holds in our system due to the relatively weak binding of E2-Ub conjugates to RING domains^10^—substrate depletion is well approximated by a simple exponential decay function (Fig. 4B). Consequently, experiments initiated at the same initial substrate concentration but conducted at different enzyme concentrations would produce a series of exponential decay curves with identical upper and lower boundaries but distinct decay rates (Fig. 4C). However, experimental data collected at varying degrader concentrations reveal clear deviations from the simple Michaelis-Menten model, suggesting that additional factors influence substrate depletion by the ternary complex (Fig. 4A,C).

In the degrader titration series, the FRET decay associated with ternary complex activity is evident only during the first 2–3 hours of the reaction, after which all samples exhibit an exponential decay toward a common final FRET value (*F*_*min*_) at a shared rate (*k*_0_), independent of degrader concentration (Fig. 4A). This late-phase decay reflects passive, E3-independent hydrolysis of the E2-Ub conjugate. The apparent loss of ternary complex enzymatic activity within the first 2-3 hours of the reaction is reminiscent of Time-Dependent Inactivation (TDI), a concept used in pharmacology to describe the gradual inactivation of cytochrome P450 enzymes^21^.

To account for this gradual loss of E3 activity, we incorporated an inactivation rate parameter (*k*_*inact*_) into the differential equation governing substrate depletion (Fig. 4D). The theoretical substrate depletion curve was then obtained by numerically solving this equation, and model parameters were determined by performing nonlinear lease-squares fitting of the theoretical curves to the data. This analysis yielded excellent agreement between the model and experiment throughout the course of the reaction for all degrader concentrations (Fig. 4E).

Although our time-dependent inactivation model contains five free parameters, several factors support the robustness of fitting procedure (*Methods*). Two of the five parameters were globally minimized across the entire degrader titration series. The three remaining parameters were strongly overdetermined by the dataset and exhibited low covariance, ensuring reliable and independent estimation (Fig. 4F). While the precise mechanism underlying gradual E3 inactivation remains unknown, the *k*_*E*3_ values extracted from the data using the time-dependent inactivation model provide a quantitative measure of the active ternary complex present at the start of each reaction.

### Extracting ternary complex characteristics from the ubiquitin transfer rates

Ternary complex formation in TPD is fundamentally a three-component binding equilibrium problem, where the binding affinities and cooperativity between the degrader, target protein, and E3 ligase dictate the overall assembly (Fig. 4A). Mathematically, this problem reduces to solving a system of three equations with three unknowns (Fig. 4B). However, the general solution is given by the root of a fifth-degree polynomial, for which no closed-form analytical solution exists^22^. While various approximations have been developed for specific limiting cases^22–25^, none are universally applicable. Nevertheless, ternary complex concentration [*EDS*] can be computed to arbitrary precision using a numerical equation solver, given known total concentrations [*E*]_*t*_, [*D*]_*t*_, [*S*]_*t*_ and three equilibrium constants 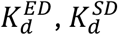 and 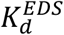.

According to Michaelis-Menten kinetics (Fig. 3B), the rate of E2-Ub consumption by the ternary complex (*k*^*EDS*^ in Fig. 4A,C) is approximately proportional to ternary complex concentration when [E2-Ub] ≪ *K*_*m*_, with the proportionality coefficient corresponding to the catalytic efficiency of the ternary complex (*k*_*cat*_/*K*_*m*_). Thus, the rate of E2-Ub depletion can be numerically calculated based on three known total concentrations ([*E*]_*t*_, [*D*]_*t*_ and [*S*]_*t*_) and four model parameters: 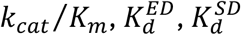 and 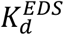. A least-squares minimization algorithm can then be applied in conjunction with this numerical solution to determine best-fit values for these four parameters from a well-sampled experimental dataset containing E2-Ub consumption rate measurements across varying ligase, substrate protein, and degrader concentrations (*Methods*).

The rate of E2-Ub depletion by the ternary complex *k*^*EDS*^ is determined experimentally by subtracting the degrader-independent background depletion rate from the *k*_*E*3_ values derived in the previous section *k*^*EDS*^([*D*]_*t*_) = *k*_*E*3_([*D*]_*t*_) − *k*_*E*3_([*D*]_*t*_ = 0) (Fig. 4D). To ensure that the four free parameters of the model are well constrained in the dataset, we performed *k*^*EDS*^ measurements at twelve different [*D*]_*t*_ concentrations for each of four distinct substrate protein concentrations [*S*]_*t*_ (Fig. 4D). Two replicate measurements were performed for each sample to evaluate the magnitude of the experimental rate measurement error. Experiments shown in Fig. 4 were performed with the BD2 (aa 333-460) construct of BRD4.

At this level of precision, the rich informational content of the dataset is evident from the distinct patterns of *k*^*EDS*^ values, which rise and fall as [*D*]_*t*_ increases, reaching maxima of varying magnitudes at different [*D*]_*t*_ values for each [*S*]_*t*_ concentration. Indeed, the standard errors of the least-squares fitting procedure, as computed from the covariance matrix, indicate strong convergence and precise parameter estimation. Further validation of the model’s stability was performed by introducing increasing levels of artificial noise into simulated datasets, confirming that the fitting procedure remains robust under realistic levels of experimental variability (*Methods*).

### Multiparametric view of TPD

The four degrader characteristics determined using the FRET assay provide a detailed, multiparametric view of ternary complex assembly and activity. As an illustrative example, we compared two well-characterized degraders, dBET1 and dBET6, both of which recruit the BRD4 substrate to CRL4^CRBN^. These heterobifunctional degraders differ only in the length of the linker connecting their two warheads: a CRBN-binding thalidomide moiety and the BRD4-binding ligand, JQ1.

The BRD4 BD2 binding affinities of dBET1 and dBET6 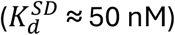, measured using our assay, are similar to each other and to the affinity reported for isolated JQ1 by isothermal titration calorimetry (ITC)^26^. In contrast, while dBET6 shows CRBN-binding affinity 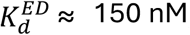 comparable to values reported for thalidomide analogs measured by ITC^27^, dBET1 exhibits nearly 10-fold tighter binding to CRBN. This suggests that the shorter linker of dBET1 may facilitate additional favorable interactions between CRBN and the degrader. For example, the formation of even a single well-oriented hydrogen bond between the JQ1 warhead and CRBN could account for a 10-fold increase in binding affinity.

Analysis of the ubiquitination rates also yields the affinity of ternary complex assembly 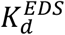 (Fig. 4A,B), which can be used to calculate assembly cooperativity defined as the ratio 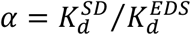^28^. For dBET6, no significant positive cooperativity is observed (α ≈ 1), consistent with crystal structures showing no direct favorable interactions between BRD4 and CRBN in the ternary complex^29^. For dBET1, the cooperativity is negative (α < 1), likely due to its shorter linker imposing steric clashes and reducing conformational entropy, thereby contributing unfavorably to ternary complex stability.

In addition to the three equilibrium constants describing ternary complex assembly, our assay also measures the value of *k*_*cat*_/*K*_*m*_, which quantifies the catalytic efficiency of the assembled ternary complex. The magnitude of this parameter can be used to evaluate whether the relative orientation of components within the ternary complex favors productive ubiquitination by positioning of the substrate lysine near the E2-Ub thioester bond.

Finally, we investigated the utility of the assay for studies of molecular glues—small molecules that induce ternary complex assembly primarily by promoting direct favorable interactions between the ligase and the substrate. For this purpose, we generated a minimal zinc finger 2 (ZF2) construct of the human protein SALL4 comprising amino acids 405-432 and investigated its ubiquitination by the CRL4^CRBN^ ligase in the presence of pomalidomide (Fig. 5A). SALL4 is a developmental transcription factor implicated in thalidomide embryopathy, and the structural basis of its recruitment to CRBN by thalidomide-related molecular glues is well characterized^30–33^. We observed that the addition of the SALL4 ZF2 construct accelerated the depletion of the FRET-active UBE2D3-Ub conjugate by CRL4^CRBN^ even when no pomalidomide was present, whereas the addition of pomalidomide further enhanced the acceleration of the FRET decay (Fig. 5B). The findings reveal that, in contrast to BRD4, the degrader-independent interaction of CRL4^CRBN^ ligase with SALL4 ZF2, described by the dissociation constant 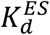, needs to be explicitly included in the system of binding equilibria (Fig. 5C). On the other hand, the affinity of pomalidomide for SALL4 is negligible 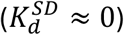 and can be ignored in the theoretical description of the system. Therefore, the molecular glue binding equilibria can be described by a different system of three equations (Fig. 5D). Another important difference between BRD4 and SALL4 ubiquitination is that both the ternary complex [*EDS*] and the pomalidomide-free CRL4^CRBN^-SALL4 complex [*ES*] catalyze ubiquitin transfer and contribute to the total, substrate-dependent FRET decay rate, *k*^*Tot*^ = *k*^*EDS*^ + *k*^*ES*^ (Fig. 5C). Experimentally *k*^*Tot*^ is determined by subtracting SALL4-independent background rate from all measured *k*_*E*3_ values: *k*^*Tot*^ = *k*_*E*3_([*D*]_*t*_, [*S*]_*t*_) − *k*_*E*3_([*D*]_*t*_ = 0, [*S*]_*t*_ = 0) (Fig. 5E).

**Figure 5.**
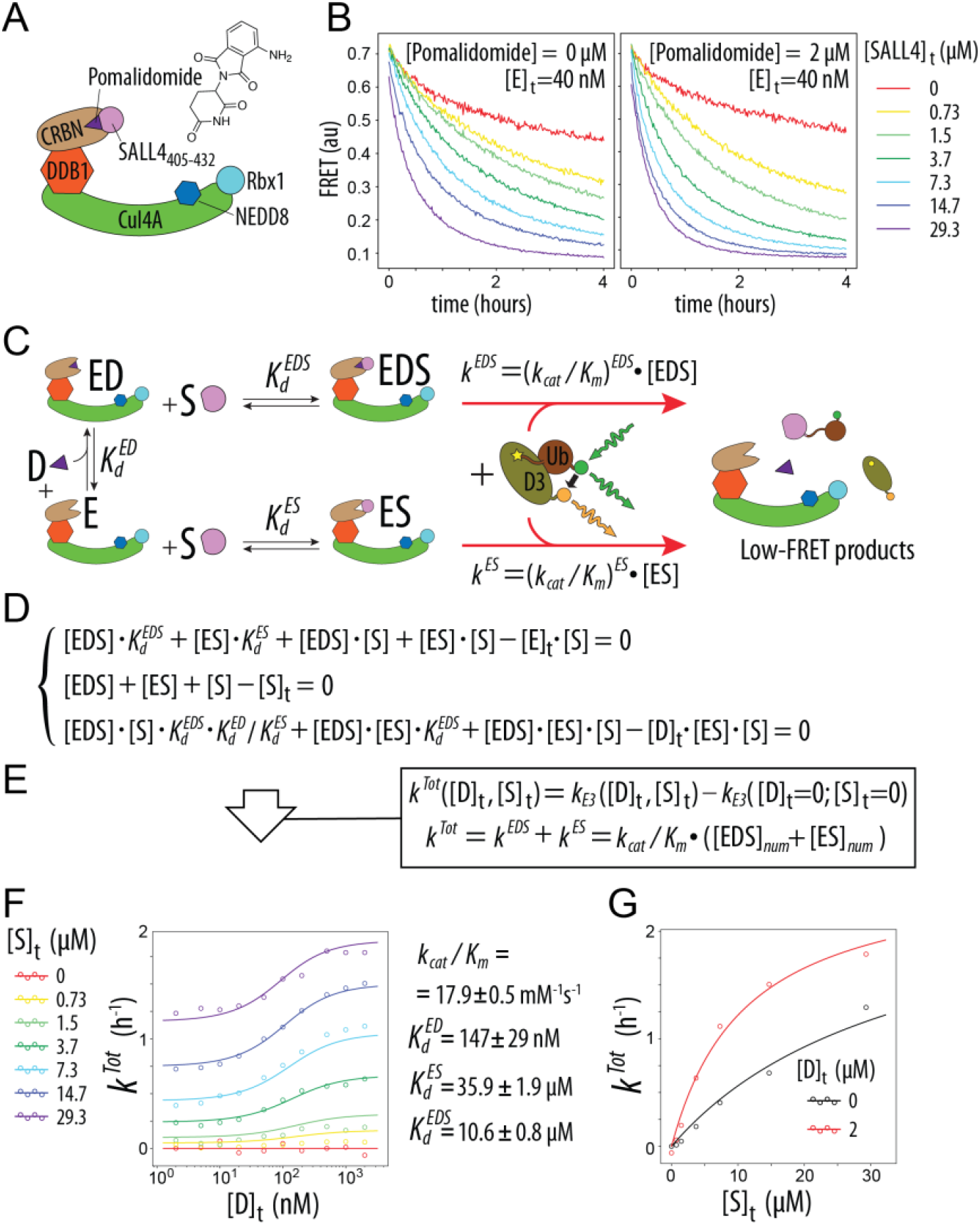
Quantitative modeling of molecular glue activity in systems with non-negligible E3-substrate affinity. (A) Recruitment of SALL4 to CRL4^CRBN^ by thalidomide and its analogues is implicated in thalidomide embryopathy. (B) FRET progress curves for ubiquitination of SALL4_405-432_ at varying substrate concentrations in the absence (left) and presence (right) of 2 µM pomalidomide. (C) Schematic of binding equilibria contributing to both molecular glue–dependent and –independent ubiquitin transfer. (D) System of mass conservation equations describing the equilibrium distribution of ED, ES, and EDS complexes. (E) Model parameters are obtained by least-squares fitting of experimental *k*^*Tot*^ values to theoretical rates obtained by numerically solving the system in panel D. (F) Experimental *k*^*Tot*^ values (circles) and model predictions (lines) across substrate and degrader titration series. (G) Plotting *k*^*Tot*^ versus SALL4_405-432_ concentration for the 0 μM and 2 μM pomalidomide datasets illustrates an approximately 3-fold increase in ligase-substrate affinity upon pomalidomide addition.

Using a computational strategy analogous to the one described in the previous section, one can determine best-fit values of model parameters from a well-sampled experimental dataset (Fig. 5E,F and *Methods*). In this analysis, we assume that the catalytic efficiencies of the [*EDS*] and [*ES*] complexes are the same, to keep the total number of free model parameters in the least-squares minimization algorithm at four: 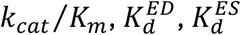 and 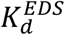. The theoretical curves calculated using best-fit parameter values are in good agreement with experimental dataset, although some systematic deviations between the theory and experiment are also noticeable (Fig. 5F). It is self-evident that the problem of model parsimony is an ever-present concern when a model with four or more free parameters is being used to describe a biochemical system containing thousands of atoms. Nevertheless, the binding affinity constants of ternary complex assembly determined by the method described here are in good general agreement with previously reported results using other biophysical techniques. Even though the absolute values of the relatively low affinities of the SALL4 ZF2 (aa construct for CRBN (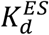 and 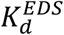) have a large margin of error, it is evident that the approximately 3-fold enhancement of the CRL4^CRBN^-SALL4 binding affinity by pomalidomide 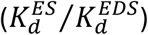 can be measured with a high degree of accuracy (Fig. 5B,G).

Collectively, our findings illustrate that the sensitivity and precision of the FRET ubiquitin transfer assay make it a powerful experimental tool for quantitative analysis of ternary complex assembly and activity.

## DISCUSSION

In this study, we demonstrate the utility of purified, FRET-active E2–Ub conjugates for quantitative analysis of ubiquitin transfer in TPD. With purified E2-Ub reagents, the E3 ligase activity can be reconstituted independently from E1-catalyzed E2-Ub synthesis, which greatly simplifies data interpretation. The high sensitivity of FRET detection allows monitoring of enzyme kinetics at physiologically relevant concentrations and in the single-step, single-turnover regime, providing a foundation for rigorous modeling of enzyme kinetics. Here, we illustrate the power of this analytical approach by measuring several key characteristics of targeted protein degraders: affinity for the target protein, affinity for the ligase, affinity of ternary complex assembly, and catalytic efficiency of the ternary complex. These parameters measured in a single experiment are in good agreement with values previously obtained using other biophysical techniques.

The methodology is highly resistant to common sources of experimental noise, as shown by the robustness of normalized FRET measurements across replicate wells and independent experiments. In addition, the assay performs well in the high-throughput format, which will facilitate screening of compound libraries in reconstituted systems containing catalytically active ligases and unmodified target proteins. Its capacity to deliver side-by-side, multiparameter evaluation of compound libraries enables efficient structure-activity relationship (SAR) exploration. Acquisition of large, high-precision datasets containing internally consistent affinity and activity parameters may facilitate the application of artificial intelligence (AI) and machine learning tools for predictive modeling. Such models could aid in correlating degrader chemical structure with ternary complex properties, an area of active interest in modern drug discovery.

One of the key strengths of our approach lies in its broad applicability to diverse ubiquitin ligases and substrates. The assay requires no engineering or modification of the E3 ligase or substrate protein, instead relying on preformed, FRET-active E2-Ub conjugates. This design avoids potential artifacts that may arise from fusion constructs or epitope tags and allows the assay to be adapted readily across different E3-substrate systems. While the assay is demonstrated here in the context of the CRL4^CRBN^ system, its modular design allows straightforward extension to other E3 ligases and ubiquitination pathways. Beyond TPD, the approach could be adapted to study natural substrate recognition, E3-substrate selectivity, and the effects of posttranslational modifications or mutations on ubiquitin transfer kinetics.

In summary, the FRET-active E2–Ub conjugates described here provide a sensitive, quantitative, high-throughput, and versatile analytical platform that enables both fundamental studies of E3 ligase function and regulation, as well as translational applications in TPD and beyond.

## MATERIALS AND METHODS

### Protein Reagents and Chemicals

Purified FRET-active E2-Ub thioester conjugates were obtained from E3 Bioscience (e3bioscience.com). Ub-R3-TRIM5α and UBE2V2 proteins were prepared as previously described^11^. Human BRD4 and SALL4 constructs were generated using standard E.coli expression and affinity purification protocols ^11^. The BRD4 constructs, BD1-BD2 (aa 44-460) and BD2 (aa 333-460) contained an N-terminal His_6_ tag. The SALL4 ZF2 construct (aa 405-432) contained an N-terminal His_10_-NusA solubility enhancement tag ^34^. Ubiquitination assays were performed on the purified fusion proteins without removing these fusion tags. Human Cullin4A and Rbx1 proteins were co-expressed in insect cells, the heterodimer purified and subsequently neddylated following published protocols ^35,36^. Human DDB1-CRBN heterodimer was similarly generated using insect-cell co-expression ^27^. dBET1 (cat. #: SML2687), dBET6 (cat. #: SML2683) and pomalidomide (cat. #: P0018) were purchased from Sigma-Aldrich.

### FRET assays

All FRET assays were performed in the buffer containing 50 mM Na phosphate pH 7.4, 100 mM NaCl and 2 mg/mL bovine serum albumin. FRET assays were performed in 384-well black low-protein-binding assay microplates. Manually dispensed test assays were performed at 40 μL final volume in Corning 3575 microplates. Low-volume, high-throughput and quantitative assays were performed at 6 μL final volume using automated dispensing of reagents into low-volume Revvity ProxiPlates (cat. #: 6059260). In 6 μL assays, degrader compounds were first dispensed as DMSO stocks at 60 nL per well using acoustic liquid handling (Echo; Beckman Coulter). The remaining components were dispensed as aqueous solutions using Formulatrix Mantis automated liquid dispenser. First, substrate protein stock and buffer were dispensed to 4 μL total volume per well to yield desired substrate protein concentration. Reconstituted E3 ligase stock was then dispensed at 1 μL per well. E3, substrate and degrader were incubated for 20 min prior to dispensing E2-Ub conjugates. E2-Ub conjugate stock was dispensed at 1 μL per well, the plates sealed with transparent film to prevent evaporation and fluorescence monitored over time using a BioTek Synergy Neo2 plate reader equipped with dual PMT detectors.

### Computational Data Analysis

Extensively commented Python scripts used for data analysis and experimental datasets are available from GitHub: https://github.com/ivanov-laboratory/TPD. Briefly, data analysis was performed in two steps: first, catalytic rates were extracted from experimental FRET decay datasets (get_rates.py), and second, these rates were analyzed as a function of total reagent concentrations to derive ternary complex characteristics (glue_fit_rates.py or protac_fit_rates.py). Each of the two steps used a solver-in-the-loop strategy: a numerical solver produced a model prediction for each trial set of parameters, and those predictions were compared to the experimental data via nonlinear least squares (scipy.optimize.least_squares). Parameter values were optimized iteratively within least_squares to minimize the sum of squared residuals between model and data.

#### FRET decay fitting

Donor and acceptor intensities were parsed from plate-reader text files; FRET was computed as the ratio of acceptor to donor fluorescence intensity signals. For each sample, the FRET decay was modeled by a time-dependent inactivation (TDI) ODE (Figure 3D), which was numerically integrated with scipy.integrate.solve_ivp. The resulting timecourse served as the theoretical model inside least_squares. Two fitting tiers were used (Fit 1 and Fit 2). Fit 1 (5-parameter):

*k*_*E*3_, *F*_*max*_ and *k*_*inact*_ were fitted per sample; *k*_0_ and *F*_*min*_ were fitted globally. This was typically applied to long acquisitions (4–6 h) on a subset of wells (or column-by-column for the whole plate). Fit 2 (reduced): For shorter traces (1–2 h) or large datasets, globally determined parameters from Fit 1 were fixed and the remaining 2 or 3 parameters were refit per sample. Options included 2-parameter, 2-plus-1 (with *k*_*inact*_ constrained linearly vs. *k*_*E*3_), or 3-parameter variants.

#### Equilibrium model fitting

Catalytic rates were fit to mass-action equilibrium models of ternary complex formation mediated by either heterobifunctional degraders (PROTACs) (Figure 4B) or molecular glues (Figure 5D). The algebraic systems enforcing mass conservation and binding equilibria were solved numerically with scipy.optimize.fsolve to obtain concentrations of the catalytically active species. These numeric predictions populated a model rate matrix that was compared to the experimental catalytic rate matrix within least_squares. Convergence of the fitting procedure and its sensitivity to experimental noise was investigated for different combinations of equilibrium constant values and reagent concentrations using scripts in the /simulate_and_analyze_rate_data folder of the GitHub repository.

## ACKNOWLEDGEMENTS

This work was supported in part by NIH R37AI136697 (D.I.); The Welch Foundation Research Grant AQ-1996-20230405 (D.I.); NIH NCI R21CA286307 (D.L.); NIH R01GM115568 (S.K.O.); R01GM128731 (S.K.O.); CPRIT RR200030 (S.K.O.). S.K.O. holds the MCC 40th Anniversary Endowed Distinguished Professorship in Oncology. We also appreciate the technical assistance provided by Dr. Daifeng Jiang at the Target Discovery Core and the Mays Cancer Center Drug Discovery Shared Resource at University of Texas Health San Antonio, supported by CPRIT Core Facility Award RP250601 and National Cancer Institute Cancer Center Support Grant P30 CA054174, respectively.

